# Use Of Canine Olfactory Detection For COVID-19 Testing Study On U.A.E. Trained Detection Dog Sensitivity

**DOI:** 10.1101/2021.01.20.427105

**Authors:** Dominique Grandjean, Dana Humaid Al Marzooqi, Clothilde Lecoq-Julien, Quentin Muzzin, Hamad Katir Al Hammadi, Guillaume Alvergnat, Kalthoom Mohammad Al Blooshi, Salah khalifa Al Mazrouei, Mohammed Saeed Alhmoudi, Faisal Musleh Al Ahbabi, Yasser Saifallah Mohammed, Nasser Mohammed Alfalasi, Noor Majed Almheiri, Sumaya Mohamed Al Blooshi, Loïc Desquilbet

## Abstract

This study aimed to evaluate the sensitivity of 21 dogs belonging to different United Arab Emirates (UAE) Ministry of Interior (MOI), trained for COVID-19 olfactory detection.

The study involved 17 explosives detection dogs, two cadaver detection dogs and two dogs with no previous detection training. Training lasted two weeks before starting the validation protocol. Sequential five and seven-cone line-ups were used with axillary sweat samples from symptomatic COVID-19 individuals (SARS-CoV-2 PCR positive) and from asymptomatic COVID-19 negative individuals (SARS-CoV-2 PCR negative). A total of 1368 trials were performed during validation, including 151 positive and 110 negative samples. Each line-up had one positive sample and at least one negative sample. The dog had to mark the positive sample, randomly positioned behind one of the cones. The dog, handler and data recorder were blinded to the positive sample location.

The calculated overall sensitivities were between 71% and 79% for three dogs, between83% and 87% for three other dogs, and equal to or higher than 90% for the remaining 15 dogs (more than two thirds of the 21 dogs).

After calculating the overall sensitivity for each dog using all line-ups, “matched” sensitivities were calculated only including line-ups containing COVID-19 positive and negative samples strictly comparable on confounding factors such as diabetes, anosmia, asthma, fever, body pain, diarrhoea, sex, hospital, method of sweat collection and sampling duration. Most of the time, the sensitivities increased after matching.

Pandemic conditions in the U.A.E., associated with the desire to use dogs as an efficient mass-pretesting tool has already led to the operational deployment of the study dogs.

Future studies will focus on comparatives fields-test results including the impact of the main COVID-19 comorbidities and other respiratory tract infections.

## INTRODUCTION

Due to the global COVID-19 pandemic, there is an increasing need for “easy to use” and rapid testing methods. COVID-19 has caused unprecedented challenges requiring a proactive and vigilant approach. At the end of December 2020 in the United Arab Emirates (U.A.E.), the average number of daily tests was around 146,000, with cumulative numbers of diagnosed COVID-19 cases and deaths of 197,000 and 645, respectively [1].

Due to potential subsequent waves of COVID-19 in many countries and the availability of large numbers of existing drug and explosives detection dogs in the U.A.E., the U.A.E Ministry of Interior decided to join the Nosaïs multicentre study in April 2020. This program is conducted by the National Veterinary School of Alfort (France) and the St Joseph University of Beirut (Lebanon) and has been established to develop the scientific approach of medical detection dogs.

Two recent studies provided evidence that detection dogs appear able to detect COVID-19 positive individuals through olfactory detection.[2, 3] In addition to investigating a new COVID-19 “testing system”, these studies also show that the “One health - One Medicine” is more important than ever as it is bringing medical doctors, veterinary surgeons, epidemiologists and dog-handlers together to share their knowledge and experience in an attempt to combat the current pandemic.

Numerous studies have suggested that dogs seem able to detect human diseases[4], such as bladder [5, 6], colon [7, 8], prostate [9–11], and liver [12] cancers, melanoma [13, 14], diabetes [15–18],epileptic fits[19], malaria [20], and bacteriological diseases [21, 22]. The high performances of the dogs in these studies encouraged further research [23]. It appeared that dogs can detect the Volatile Organic Compounds (VOCs) generated during the conditions but the specificity of this induced odour to the microorganism was unclear. Several studies explored the VOCs induced by infectious processes but these focussed on the disease-related inflammatory reaction or oxidative stress, without looking at the specificities of the involved pathogen [24–28]. In 2014, Aksenov et al. [29] demonstrated that the VOCs produced by cell-cultures infected by three different types of Influenza viruses (H9N2, H6N2 and H1N1) were specific to each virus. More recently, Abd el Qader et al. [30] and Schivo et al. [31] reached the same conclusion with different rhinoviruses. Therefore, there is a high probability that coronaviruses follow the same rule and that SARS-CoV-2 generates specific VOCs [32] but this is yet to be proven.

Training dogs for acute medical detection is not easy, as the whole process requires a large number of high-quality samples (both positive and negative samples in the case of COVID-19). But once correctly trained, dogs can be used in real-time to detect whether or not individuals are infected by an active SARS-CoV-2 virus.

To provide evidence that dogs can detect COVID-19 positive individuals, the study protocol must follow recommendations to prevent biases and over-interpretation of the results.[33–35] These recommendations include ensuring the dog handler is unaware of the individual’s disease status when presenting to the dog, ensuring the dog is presented one sample no more than once during the training and the validation sessions, ensuring control samples are comparable to positive samples except for disease status (to avoid confounding bias), and randomising sample positions in the line-up when used. This study aimed to estimate the individual sensitivity of dogs trained to detect COVID-19 positive individuals.

## MATERIALS AND METHODS

This study was conducted using the guidelines written by the Nosaïs program team of Alfort National Veterinary School (France) [3], and was conducted in strict accordance with the ethical principles in the Declaration of Helsinki (60^th^ General Assembly of the AMM, Seoul-Korea, October 2013).

### Samples

COVID-19 positive and negative axillary sweat samples used for the training and the validation sessions were collected by doctors and nurses in assigned U.A.E government hospitals (listed in Table 1) who were trained not to contaminate the samples with their own odours. The reasons for choosing sweat, the sampling site and method, and biological safety measures have been described previously by Grandjean et al. [3].

**Table 1.** List of the U.A.E. MOHAP hospitals involved in sample collection for the training and validation sessions.

|  |
| --- |
| AL QASSIMI HOSPITAL |
| SAQR HOSPITAL |
| UM ALQUWAIN HOSPITAL |
| KALBA HOSPITAL |
| FUDAIRAH HOSPITAL |
| SHARJAH FIELD HOSPITAL |
| AL QUASSIMI WOMEN AND CHILD HOSPITAL |
| MASAFI HOSPITAL |
| OBAIDALLAH HOSPITAL |
| KHORFKHAN HOSPITAL |
| RASHID HOSPITAL |
| SKML HOSPITAL |

Patients presenting to one of the participating hospitals with COVID-19 clinical symptoms (such as fever, cough, throat pain, malaise or body pain), and having a positive Reverse Transcription Polymerase Chain Reaction (RT-PCR) or PCR test for SARS-CoV-2 were included as positive individuals. Patients from the participating hospitals with a negative COVID-19 PCR test result were included as negative individuals. All individuals meeting these inclusion criteria were asked if they were willing to participate in the study and signed an individual informed consent form approved by the national ethics committee.

Sweat samples were collected using Getxent® inert polymers tubes positioned in direct contact with the skin (occasionally over the first layer of clothes) in the patient’s axilla for up to 20 minutes. The tubes were stored in tinted glass jars (to prevent UV ray damage) placed in individual biohazard bags. The bags were marked with the patients’ individual anonymous code and the RT-PCR results. Positive and negative samples were separated and stored in a refrigerator. No samples were screened for other human coronaviruses like beta coronaviruses HCoV-OC43 or alpha coronavirus HCoV-229E.

### Individual data collection

Medical staff recorded demographic and medical data about the included individuals, strictly respecting local regulations. These data included: age, sex, weight, and the presence or absence of hypertension, diabetes, fever, dry cough, body pain, sore throat, diarrhoea, asthma, and anosmia.

### Sample transportations

Once collected, samples were transported from the hospital to the Dubaï Police K9 Training Centre once a day by a dedicated Ministry of the Interior driver. All individually packaged samples were transported in two-compartment medical coolers with icepacks ensuring the positive and negative samples remained separate.

### Sample storage

Samples were stored in boxes (keeping positive and negatives samples separate) in a refrigerator at 4°C until they were used for the dog training or validation sessions. They were never manipulated without disposable surgical gloves to prevent any odour contamination.

### Canine resources

A total of 21 dogs participated in the study belonging to the following U.A.E institutions: Dubai Police, Fujairah Police, Ajman Police, Ras Al Khaimah Police, Sharjah Police, U.A.E Land forces, and U.A.E Federal Customs. They were explosives detection dogs (n=17), cadaver detection dogs (n=2), and “green” dogs (i.e. dogs without previous olfactory detection training) (Table 2). Explosives detection dogs were chosen because they are already trained to detect 20 to 30 different explosives odours so the potential COVID-19 specific odour would be easily memorised and generalised. Cadaver detection dogs are accustomed to olfactory detection and their training does not interfere with a new odour being imprinted. “Green” dogs begin training with a short period of nose work.

**Table 2:** Dogs involved in the study

| Name | Gender | Breed | Age | Organisation | Speciality |
| --- | --- | --- | --- | --- | --- |
| ACE | Male | Malinois | 1 | Dubai K9 | Green Dog |
| ALLO | Male | Malinois | 3 | Fujairah Federal K9 | Explosives |
| AXEL | Male | German Shepherd | 6 | UAE Land Forces | Explosives |
| BEN3 | Male | Labrador | 6 | Dubai K9 | Cadaver |
| BOLT | Male | Malinois | 1 | Dubai K9 | Green Dog |
| BOLTON | Male | German Shepherd | 2 | Sharjah Federal K9 | Explosives |
| BRAKEN | Male | Cocker Spaniel | 2 | Dubai K9 | Explosives |
| BREN | Male | Malinois | 3 | Ajman Federal K9 | Explosives |
| CODY | Male | Cocker Spaniel | 2 | Dubai K9 | Explosives |
| CUBA | Male | Dutch Shepherd | 4 | Federal Customs | Explosives |
| FLASH | Male | Cocker Spaniel | 3 | Dubai K9 | Explosives |
| FUDU | Male | German Shepherd | 3 | Ajman Federal K9 | Explosives |
| KIRA | Female | Malinois | 4 | Federal Customs | Explosives |
| MILO | Male | Cocker Spaniel | 3 | Dubai Customs | Explosives |
| NEJAN | Male | Malinois | 4 | Fujairah Federal K9 | Explosives |
| NERO | Male | Cocker Spaniel | 2 | Dubai Customs | Explosives |
| NOX | Male | Malinois | 6 | UAE Land Forces | Explosives |
| PABLO | Male | Cocker Spaniel | 2 | Dubai K9 | Explosives |
| RAMOS | Male | German Shepherd | 5 | Sharjah Federal K9 | Cadaver |
| RICKY | Male | Malinois | 6 | UAE Land Forces | Explosives |
| SPIKE | Male | Pointer | 2 | Sharjah Federal K9 | Explosives |

### Biological safety features

Despite a few publications suggesting that infected dogs may transmit the virus back to humans [36], the World Organisation for Animal Health attests that, for dogs, infection susceptibility is low and there is no transmission risk[37]. Based on these facts and in addition to the biological safety measures described for the sweat samples, measures were implemented including dog handlers and assistants wearing masks and disposable gloves, no petting with bare-hands, and no kissing the dog or allowing the dog to lick. The training and validation sessions used five and seven-cone line-ups (see below) which prevented the dog contacting the samples (Figure 1). Furthermore, for safety reasons, samples were not used for training or validation sessions within 24 hours of collection.

**Figure 1.**
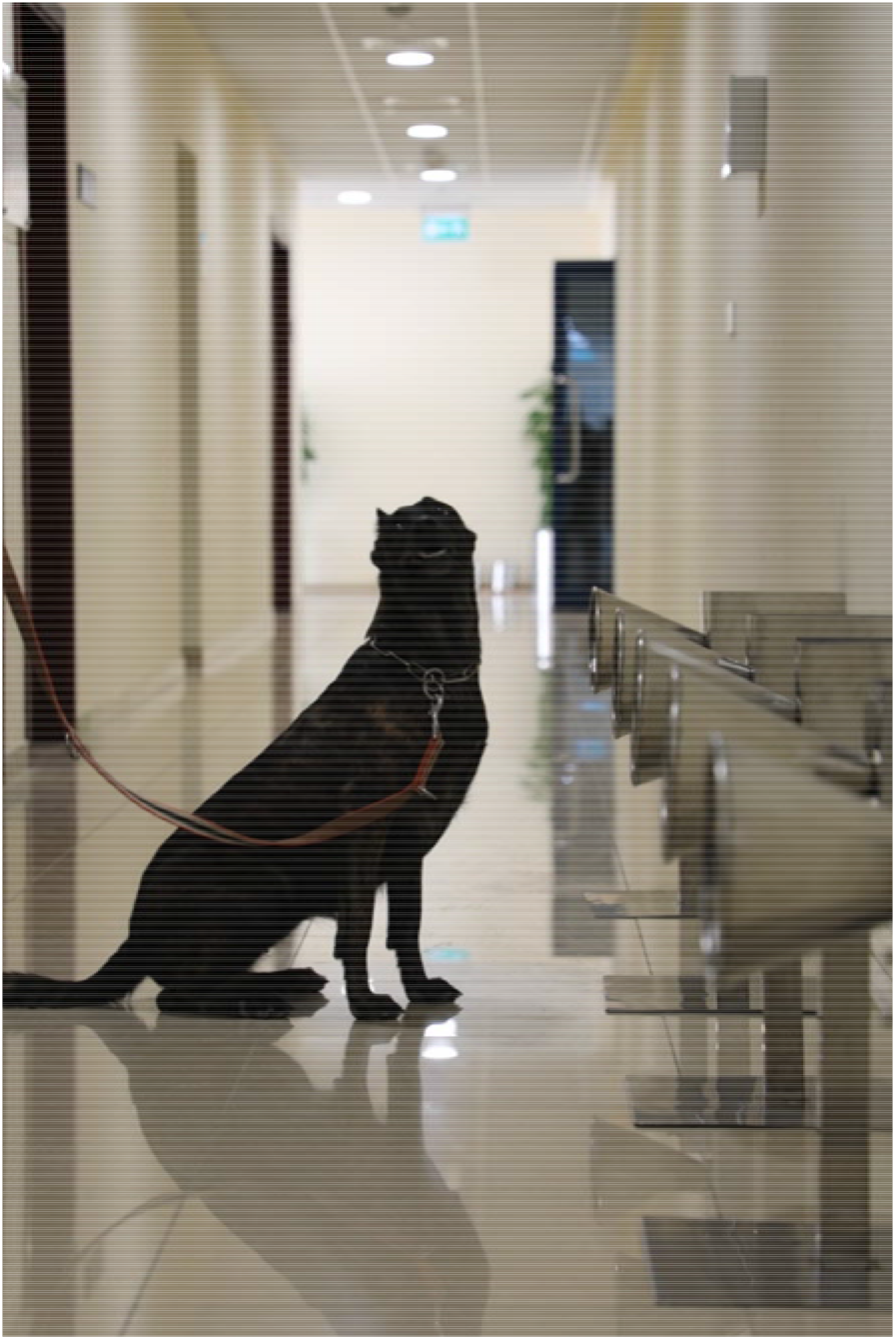
A dog positively marking a cone on a 5-cone line-up; dogs have no contact with samples

### Training protocol

As previously described,[3] dogs were trained to work on an olfactory cone line-up (Figure 2) and mark the cones containing the COVID-19 positive sample by sitting in front of it. The training sessions started on June 6^th^, 2020 and took place at the Dubai Police K9 Training Centre. All dogs were trained for eight hours daily over a two week period. The training process followed a 4-step procedure:: (step 1) learning line-up work, (step 2) memorising the COVID-19 sample odour (positive samples and empty cones in the line-up), (step 3) introducing mocks (positive samples and empty inert polymers tubes in the line-up), and (step 4) introducing negative samples without mocks in the line-up (around 30% of samples in the line-ups were COVID-19 positive). Steps 1 and 2 were carried out in week one and steps 3 (1 day) and 4 (5 days) were carried out in week two. At the end of week two, the handlers judged whether their dog was trained and ready for the validation process. Table 3 shows the number of samples (either COVID-19 positive or negative) sniffed by the dogs during step 4 of training and the number correctly marked (i.e. the dog only marked COVID-19 positive samples).

**Table 3.** Performance of the 21 dogs during the last step of the training process (step 4).

| Dog Name | Number of samples*<br>sniffed by the dog | Correct indications**, n (%) |
| --- | --- | --- |
| Ace | 81 | 72 (89) |
| Allo | 77 | 77 (100) |
| Axel | 106 | 83 (78) |
| Ben3 | 85 | 80 (94) |
| Bolt | 71 | 67 (94) |
| Bolton | 55 | 51 (93) |
| Braken | 78 | 77 (99) |
| Bren | 75 | 58 (77) |
| Cody | 89 | 88 (99) |
| Cuba | 95 | 89 (94) |
| Flash | 84 | 84 (100) |
| Fudu | 96 | 67 (70) |
| Kira | 86 | 86 (100) |
| Milo | 80 | 74 (93) |
| Nejan | 88 | 85 (97) |
| Nero | 85 | 85 (100) |
| Nox | 97 | 83 (86) |
| Pablo | 87 | 87 (100) |
| Ramos | 99 | 92 (93) |
| Ricky | 86 | 83 (97) |
| Spike | 86 | 68 (79) |
| * Samples were either COVID-19 negative or positive; ** The dog did not mark a COVID-19 negative sample and marked a COVID-19 positive sample. |  |  |

**Figure 2.**
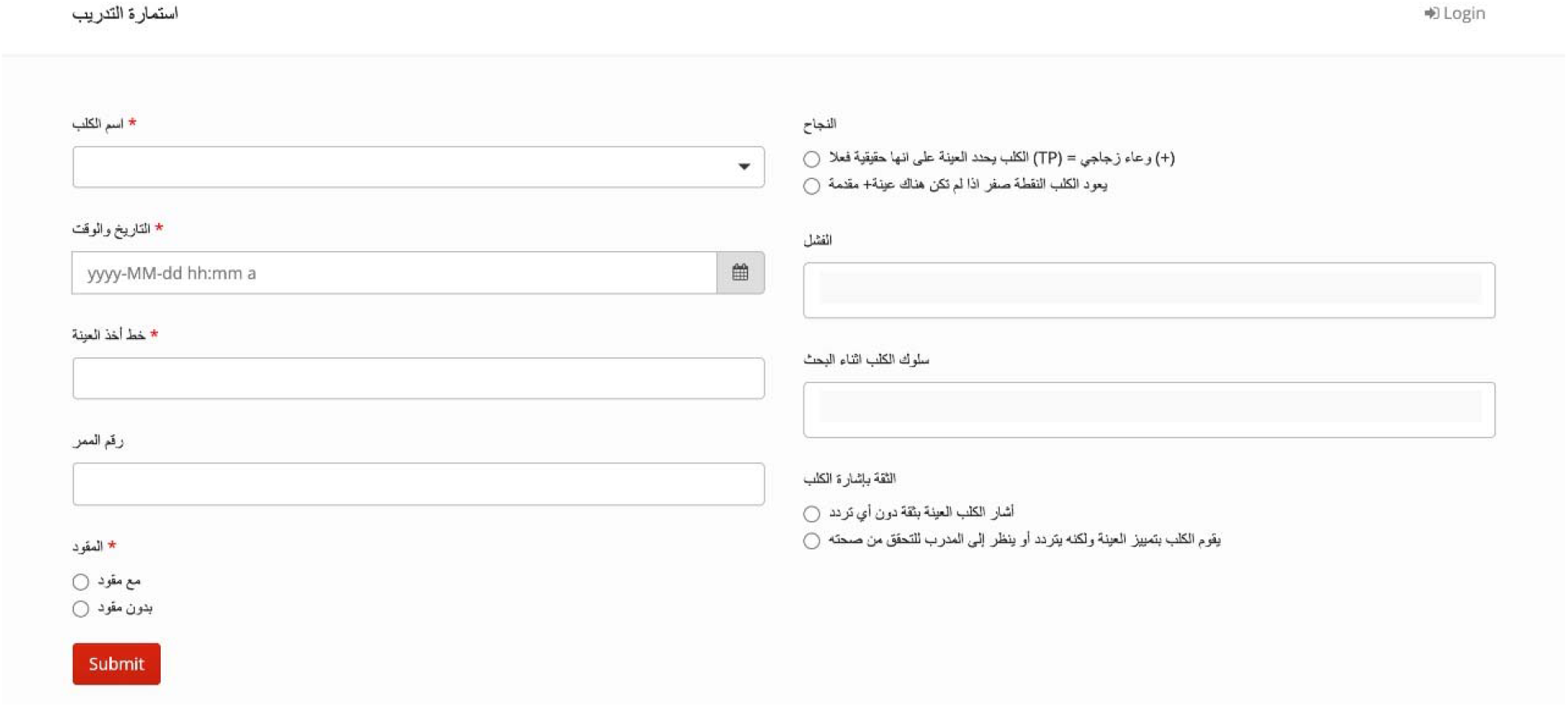
Dedicated computer software used to register data obtained during training and validation

### Validation protocol

The validation sessions took place in Dubaï Police K9 Training Centre. No samples used for training were used for validation. Most of the time, 5-cone olfactory line-ups were used. Seven-cone line-ups were used at the end of validation when dog handlers considered their dog ready for deployment in the field. There was always only one COVID-19 positive sample in the line-up, with the remaining cones containing COVID-19 negative sample(s) (at least one in the line-up) or being empty (i.e. without sampling material). The COVID-19 positive and negative sample locations in the line-up were randomly assigned and samples were placed by one dedicated person who remained in the room. The data recorder collected data in the room using dedicated software developed by Dubaï Police (Figure 2) and was blinded to the COVID-19 positive and negative sample locations. Once the samples were placed behind the cones, the handler and the dog entered the room, and the dog was asked to sniff each cone, one by one. There was no visual contact between the dedicated sample placement person and either the dog or the dog handler. If the dog marked a cone, the handler asked the dog to resume the task for the remaining cones in the line-up (i.e. sequential line-up). The dog handler considered a cone marked when the dog sat, stopped or barked. Each time the dog marked (whether correctly or incorrectly), the dog handler gave a reward and the data recorder recorded it. Once all the cones in the line-up had been sniffed, the dog handler and data recorder were informed of the COVID-19 positive sample location. The handler then knew if the dog had failed to mark the COVID-19 positive sample too many times for corrective measures (which did not happen). Only the COVID-19 positive sample results (i.e. presence or absence of marking) were recorded for statistical analyses.

### Statistical analysis

Since only the COVID-19 positive sample results were available, sensitivities but not specificities were calculated. All line-ups including a COVID-19 positive sample which had been sniffed in a previous line-up were removed from the sensitivity calculations for this dog. Overall sensitivities and “matched” sensitivities were calculated for each dog.

The overall sensitivity was calculated by dividing the number of correctly marked COVID-19 positive samples by the total number of COVID-19 positive samples it had sniffed (one per line-up). The 95% confidence interval (CI) was calculated using Jeffreys’ method [38].

The comparability criterion (i.e. positive samples are comparable to negative samples in the line-up in all ways, except for the disease status) is a key criterion in detection dog studies to prevent confounding bias occurring[34, 39]. To provide evidence that confounding bias did not elevate the overall sensitivity, “matched” sensitivities were calculated for line-ups in which the COVID-19 positive and negative sample(s) were matched for health conditions, age, sex, hospital, sampling duration, or the sweat collection method (underarm or over clothes). The potentially confounding health conditions used for matching included: diabetes, anosmia, asthma, fever, body pain, and diarrhoea. Hypertension, dry cough, and sore throat were not matching variables since they were not thought to be potential attractors for dogs. The line-up was considered “matched” for one health condition if the health condition of the COVID-19 positive patient in the line-up was the same (either present or absent) as all the COVID-19 negative patients in that line-up. If health condition data were missing for any line-up samples, the line-up was not considered “matched” for this condition. The same matching method was used for sweat collection method. If sampling duration was equal to 20 minutes for all the line-up samples, the line-up was considered “matched”. To be “matched” for age, the difference between the age of the COVID-19 positive and negative patients in the line-up was ≤15 years. Matching was performed one characteristic at a time.

Binary and qualitative variables were presented as numbers and proportions, and quantitative variables (age and weight) were presented as medians and interquartile ranges. The Spearman rank correlation coefficient was calculated to quantify the association between the dogs’ performance in step 4 of training and the overall sensitivities calculated during validation. Statistical analyses were performed using SAS® University Edition (SAS Institute Inc., Cary, NC, USA).

## RESULTS

A total of 151 COVID-19 positive patients and 110 COVID-19 negative patients were recruited, producing 261 sweat samples used only in the validation line-ups. The 151 COVID-19 positive patients were recruited mainly from Sharjah Field and Al Qasimi hospitals, and the 110 COVID-19 negative patients were recruited mainly from Al Qasimi, Rashid, and Saqr hospitals (Table 4). The proportion of females was higher among COVID-19 negative patients (36%) compared with COVID-19 positive patients (6%). The median age and weight were similar between COVID-19 negative and positive patients (40 vs 36 years and 75 vs 70 kg, respectively). Similarly, the proportions of patients presenting fever and body pain were similar between COVID-19 negative and positive patients (10% vs 10% and 20% vs 17%, respectively). Hypertension and diabetes were more frequent in COVID-19 negative patients (36% and 30%, respectively) than in COVID-19 positive patients (15% and 14%, respectively). Sore throat, diarrhoea, asthma, and anosmia were present in ≤2% of the 261 study participants. Sweat was collected over clothes (as opposed to direct skin contact) in 14% of COVID-19 negative patients and 3% of COVID-19 positive patients (Table 5). For 84% of COVID-19 negative patients and 85% of COVID-19 positive patients, the sampling material remained in contact with the skin for 20 minutes. The median number of dogs sniffing one COVID-19 negative sample was five dogs (interquartile range, four to seven dogs; range, one to 19 dogs); it was also five dogs (interquartile range, four to five dogs; range, one to 18 dogs) for COVID-19 positive samples.

**Table 4.** Baseline characteristics of the study patients

| Variables | Overall (n=261) | PCR-negative (n=110) | PCR-positive (n=151) |
| --- | --- | --- | --- |
| Female, n (%) | 49 (19) | 40 (36) | 9 (6) |
| Hospital |  |  |  |
| Al Qasimi Hospital, n (%) | 71 (27) | 55 (50) | 16 (11) |
| Fujairah Hospital, n (%) | 5 (2) | 0 | 5 (3) |
| Kalba Hospital, n (%) | 10 (4) | 6 (5) | 4 (3) |
| Rashid Hospital, n (%) | 29 (11) | 24 (22) | 5 (3) |
| Saqr Hospital, n (%) | 25 (10) | 19 (17) | 6 (4) |
| Sharjah Field Hospital, n (%) | 110 (42) | 1 (1) | 109 (72) |
| Umm Al Quwain Hospital, n (%) | 11 (4) | 5 (5) | 6 (4) |
| Age (years)*,<sup>a</sup> | 38 [30-53] | 40 [31-58] | 36 [30-50] |
| Weight (kg)*,<sup>b</sup> | 71 [61-81] | 75 [62-85] | 70 [61-80] |
| Data on health conditions available, n (%) | 225 (86) | 86 (78) | 139 (92) |
| Hypertension, n (%) | 52 (23) | 31 (36) | 21 (15) |
| Diabetes, n (%) | 45 (20) | 26 (30) | 19 (14) |
| Fever, n (%) | 41 (18) | 17 (20) | 24 (17) |
| Dry cough, n (%) | 35 (16) | 10 (12) | 25 (18) |
| Body pain, n (%) | 23 (10) | 9 (10) | 14 (10) |
| Sore throat, n (%) | 4 (2) | 0 | 4 (3) |
| Diarrhoea, n (%) | 4 (2) | 2 (2) | 2 (1) |
| Asthma, n (%) | 3 (1) | 3 (3) | 0 |
| Anosmia, n (%) | 2 (1) | 1 (1) | 1 (1) |
| * Median [interquartile range]; <sup>a</sup>, missing data on age for one COVID-19 negative patient and for seven COVID-19 positive patients; <sup>b</sup>, missing data on weight for 27 COVID-19 negative patients and for 13 COVID-19 positive patients. |  |  |  |

**Table 5.**
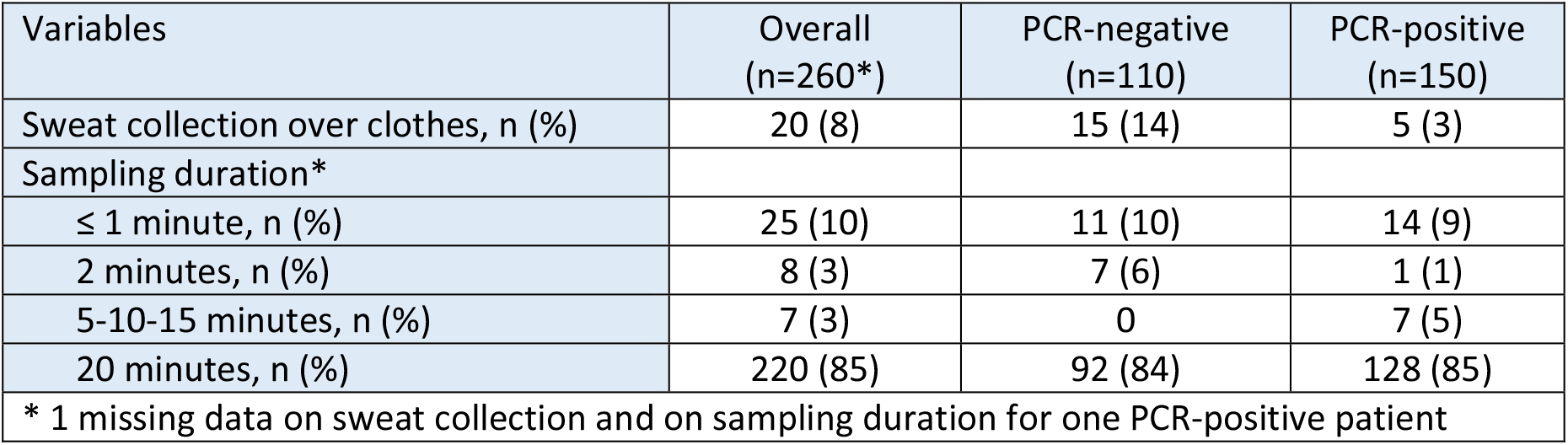
Characteristics for samples sniffed by at least one dog.

| Variables | Overall<br>(n=260*) | PCR-negative<br>(n=110) | PCR-positive<br>(n=150) |
| --- | --- | --- | --- |
| Sweat collection over clothes, n (%) | 20 (8) | 15 (14) | 5 (3) |
| Sampling duration* |  |  |  |
| ≤ 1 minute, n (%) | 25 (10) | 11 (10) | 14 (9) |
| 2 minutes, n (%) | 8 (3) | 7 (6) | 1 (1) |
| 5-10-15 minutes, n (%) | 7 (3) | 0 | 7 (5) |
| 20 minutes, n (%) | 220 (85) | 92 (84) | 128 (85) |
| * 1 missing data on sweat collection and on sampling duration for one PCR-positive patient |  |  |  |

A total of 1786 trials were performed during training, and 1368 during validation. The minimum number of line-ups performed by the dogs was 15 (Bolton), and the maximum number was 60 (Cuba; Table 6), with a median number of 35 line-ups (interquartile range, 27 to 48 line-ups). The median number of samples (either COVID-19 positive or negative) per line-up was two for 13 dogs, three for seven dogs, and four for one dog. Seven-cone line-ups (as opposed to 5-cone) were used for nine dogs at the end of the validation. When considering all line-ups performed by each dog, the overall sensitivity ranged from 71% (Fudu) to 100% (Nox, Allo, Ramos, Cody, and Bolt) (Table 6). The sensitivity was >80% in 18 of the 21 study dogs, and ≥ 90% in 15 of the 21 dogs. The 95% confidence interval lower bound was >80% in 12 of the 21 dogs. The dogs’ performances during step 4 of training were correlated to overall sensitivities during validation (Spearman rank correlation r = 0.62; p < 0.01; Figure 3).

**Table 6.** Line-up characteristics and overall sensitivities estimated for the 21 dogs during the validation process

| Dog | Number of line-ups* | Median [interquartile range] number of samples in line-ups (number of 7-cone line-ups**) | Number of correct identifications*** | Overall Se (95% CI) |
| --- | --- | --- | --- | --- |
| Ace | 43 | 3 [2 to 4] (0) | 37 | 86% (73% to 94%) |
| Allo | 27 | 2 [2 to 3] (0) | 27 | 100% (91% to 100%) |
| Axel | 23 | 2 [2 to 2] (0) | 19 | 83% (64% to 94%) |
| Ben3 | 42 | 3 [2 to 4] (0) | 38 | 90% (79% to 97%) |
| Bolt | 54 | 3 [2 to 4] (1) | 54 | 100% (95% to 100%) |
| Bolton | 15 | 2 [2 to 3] (0) | 14 | 93% (73% to 99%) |
| Braken | 23 | 2 [2 to 4] (1) | 22 | 96% (81% to 100%) |
| Bren | 29 | 2 [2 to 3] (0) | 23 | 79% (62% to 91%) |
| Cody | 39 | 2 [2 to 4] (1) | 39 | 100% (94% to 100%) |
| Cuba | 60 | 4 [3 to 4] (3) | 54 | 90% (81% to 96%) |
| Flash | 50 | 2 [2 to 4] (5) | 47 | 94% (85% to 98%) |
| Fudu | 21 | 2 [2 to 2] (0) | 15 | 71% (50% to 87%) |
| Kira | 52 | 3 [2 to 4] (3) | 48 | 92% (83% to 97%) |
| Milo | 54 | 3 [3 to 4] (3) | 47 | 87% (76% to 94%) |
| Nejan | 32 | 2 [2 to 3] (0) | 29 | 91% (77% to 97%) |
| Nero | 48 | 3 [2 to 4] (4) | 47 | 98% (91% to 100%) |
| Nox | 25 | 2 [2 to 3] (0) | 25 | 100% (91% to 100%) |
| Pablo | 35 | 3 [2 to 4] (1) | 34 | 97% (87% to 100%) |
| Ramos | 37 | 2 [2 to 3] (0) | 37 | 100% (93% to 100%) |
| Ricky | 30 | 2 [2 to 3] (0) | 29 | 97% (85% to 100%) |
| Spike | 30 | 2 [2 to 4] (0) | 23 | 77% (60% to 89%) |
| * Each 5-cone or 7-cone line-up contained one COVID-19 positive sample only, and at least one COVID-19 negative sample. ** All 7-cone line-ups contained seven samples (one COVID-19 positive and six COVID-19 negative samples). *** Number of times the dog marked the COVID-19 positive sample in the line-up. Se, sensitivity; CI, confidence interval. |  |  |  |  |

**Figure 3.**
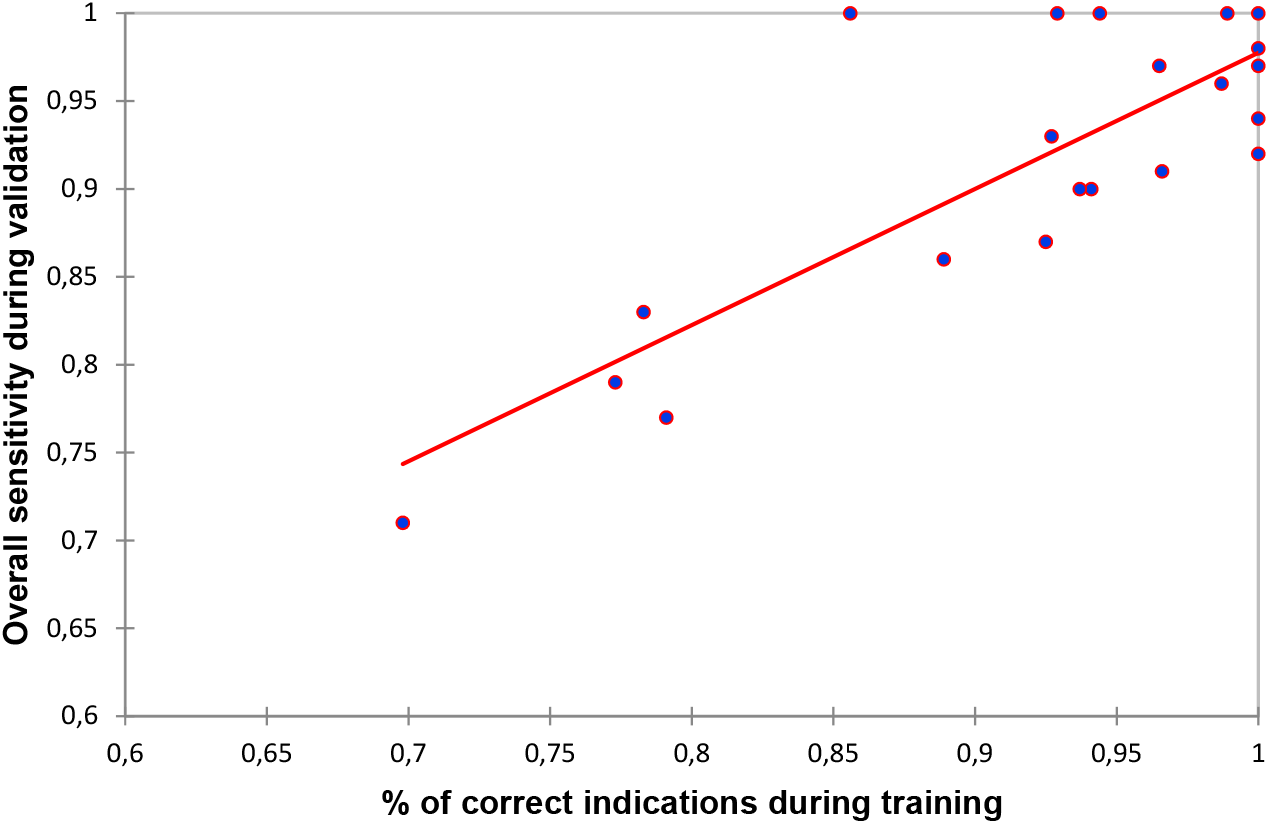
Scatter plot of the correlation between the performances of the 21 dogs during step 4 of training and overall sensitivities during validation. Spearman rank correlation r = 0.62 (p < 0.01).

Table 7 presents all “matched” sensitivities calculated for each dog using the matched line-ups. For instance, Fudu had six line-ups which were “matched” for diabetes. Fudu marked five of these six COVID-19 positive samples, leading to a diabetes “matched” sensitivity of 83%. Table 7 also highlights that in most cases, “matched” sensitivities are equal to or higher than the overall sensitivity calculated before matching. The matching characteristics resulting in decreased sensitivity were anosmia and asthma, with sensitivity only decreasing in eight of the 21 dogs.

**Table 7.** “Matched” sensitivities for the 21 dogs calculated using line-ups where the COVID-19 positive and negative samples were matched for health conditions, sex, age, hospital, sweat collection, or sampling duration.

| Dog (overall Se) | Matching variables |  |  |  |  |  |  |  |  |  |  |
| --- | --- | --- | --- | --- | --- | --- | --- | --- | --- | --- | --- |
|  | Diabetes | Asthma | Anosmia | Fever | Body pain | Diarrhoea | Sex | Age | Hospital | Sweat collection | Sampling duration |
| Ace (86) | 5/9 (56) | 16/20 (80) | 15/19 (79) | 14/17 (82) | 15/18 (83) | 16/20 (80) | 18/20 (90) | 10/11 (91) | 1/2 (50) | 37/43 (86) | 34/38 (89) |
| Allo (100) | 16/16 (100) | 18/18 (100) | 18/18 (100) | 10/10 (100) | 10/10 (100) | 18/18 (100) | 15/15 (100) | 8/8 (100) | 4/4 (100) | 27/27 (100) | 24/24 (100) |
| Axel (83) | 4/7 (57) | 9/12 (75) | 8/11 (72) | 3/4 (75) | 6/7 (86) | 9/11 (83%) | 11/12 (92) | 6/6 (100) | 7/8 (88) | 19/23 (83) | 17/21 (81) |
| Ben3 (90) | 7/9 (78) | 14/16 (88) | 13/15 (87) | 12/14 (86) | 13/14 (93) | 14/16 (88) | 18/20 (90) | 15/15 (100) | 1/1 (100) | 37/41 (90) | 30/34 (88) |
| Bolt (100) | 9/9 (100) | 21/21 (100) | 22/22 (100) | 21/21 (100) | 21/21 (100) | 23/23 (100) | 25/25 (100) | 19/19 (100) | 2/2 (100) | 53/53 (100) | 42/42 (100) |
| Bolton (93) | 5/5 (100) | 10/11 (91) | 10/11 (91) | 6/6 (100) | 8/8 (100) | 10/11 (91) | 10/10 (100) | 5/5 (100) | 3/4 (75) | 14/15 (93) | 14/15 (93) |
| Braken (96) | 4/4 (100) | 8/9 (89) | 9/10 (90) | 5/5 (100) | 3/4 (75) | 8/9 (89) | 14/14 (100) | 8/8 (100) | 4/5 (80) | 21/22 (95) | 18/18 (100) |
| Bren (79) | 5/7 (71) | 14/18 (78) | 13/17 (76) | 4/5 (80) | 12/13 (92) | 14/17 (82) | 14/16 (88) | 5/6 (83) | 7/8 (88) | 23/29 (79) | 19/25 (76) |
| Cody (100) | 9/9 (100) | 19/19 (100) | 20/20 (100) | 15/15 (100) | 12/12 (100) | 20/20 (100) | 15/15 (100) | 14/14 (100) | 7/7 (100) | 39/39 (100) | 33/33 (100) |
| Cuba (90) | 10/10 (100) | 27/27 (100) | 29/30 (97) | 21/22 (95) | 25/26 (96) | 30/31 (97) | 23/24 (96) | 19/20 (95) | 2/2 (100) | 53/59 (90) | 45/50 (90) |
| Flash (94) | 10/10 (100) | 22/22 (100) | 24/24 (100) | 15/15 (100) | 13/13 (100) | 22/22 (100) | 24/24 (100) | 18/20 (90) | 9/9 (100) | 46/49 (94) | 38/41 (93) |
| Fudu (71) | 5/6 (83) | 9/11 (83) | 8/10 (80) | 3/4 (75) | 5/6 (83) | 9/10 (90) | 9/10 (90) | 3/6 (50) | 8/8 (100) | 15/21 (71) | 13/19 (68) |
| Kira (92) | 11/11 (100) | 24/25 (96) | 25/27 (93) | 12/14 (86) | 16/18 (89) | 25/27 (93) | 22/23 (96) | 18/20 (90) | 7/8 (88) | 47/51 (92) | 42/45 (93) |
| Milo (87) | 8/11 (73) | 18/21 (86) | 18/22 (82) | 18/21 (86) | 17/20 (85) | 19/23 (83) | 20/23 (87) | 18/20 (90) | 2/2 (100) | 46/53 (87) | 42/47 (89) |
| Nejan (91) | 12/15 (80) | 19/22 (86) | 19/22 (86) | 9/9 (100) | 12/12 (100) | 19/22 (86) | 18/18 (100) | 6/7 (86) | 7/7 (100) | 29/32 (91) | 26/29 (90) |
| Nero (98) | 10/10 (100) | 20/20 (100) | 21/21 (100) | 14/14 (100) | 12/12 (100) | 21/21 (100) | 19/19 (100) | 20/20 (100) | 7/7 (100) | 46/47 (98) | 40/41 (98) |
| Nox (100) | 5/5 (100) | 14/14 (100) | 13/13 (100) | 4/4 (100) | 11/11 (100) | 13/13 (100) | 14/14 (100) | 6/6 (100) | 7/7 (100) | 25/25 (100) | 22/22 (100) |
| Pablo (97) | 10/10 (100) | 18/18 (100) | 19/19 (100) | 13/13 (100) | 10/10 (100) | 19/19 (100) | 17/17 (100) | 12/12 (100) | 7/8 (88) | 33/34 (97) | 30/31 (97) |
| Ramos (100) | 19/19 (100) | 26/26 (100) | 26/26 (100) | 14/14 (100) | 14/14 (100) | 26/26 (100) | 21/21 (100) | 13/13 (100) | 10/10 (100) | 37/17 (100) | 35/35 (100) |
| Ricky (97) | 16/16 (100) | 21/21 (100) | 21/21 (100) | 9/9 (100) | 12/12 (100) | 21/21 (100) | 16/17 (94) | 9/9 (100) | 5/5 (100) | 29/30 (97) | 27/27 (100) |
| Spike (77) | 5/8 (63) | 15/19 (79) | 14/18 (78) | 4/4 (100) | 12/12 (100) | 14/18 (78) | 13/16 (81) | 5/7 (71) | 7/9 (78) | 23/30 (100) | 20/26 (77) |
| Numbers are expressed as n/N (“matched” Se), where n is the number of correct identifications, N is the total number of line-ups containing one COVID-19 positive sample and at least one COVID-19 negative sample matched for the matching variable, and Se the sensitivity expressed as a percentage. Grey cells indicate when the “matched” sensitivity calculated in matched line-ups was equal to or higher than the overall sensitivity calculated in all line-ups (see Table 4). |  |  |  |  |  |  |  |  |  |  |  |

## DISCUSSION

During this validation process, a total of 261 samples (110 COVID-19 negative and 151 COVID-19 positive samples) were sniffed by the 21 study dogs. The estimated overall sensitivities were between 71% and 79% for three dogs, between 83% and 87% for three other dogs, and ≥90% for the remaining 15 dogs (representing 71% of the 21 dogs). These results may appear heterogeneous but considering other COVID-19 detection methods require a sensitivity ≥90% before implementation, more than two thirds of the 21 trained dogs could be selected to start work.

Four of the 21 dogs did not reach a sensitivity of 85% (Axel, Bren, Fudu, and Spike). Interestingly, these dogs also had the lowest performances during step 4 of training. The significant correlation between the training and validation performances (Figure 4) could arise from chance, confounding bias (i.e. no causal association), or a true causal association. If confounding bias is causing the significant correlation, it means there is at least one intrinsic or extrinsic characteristic of the dyad (the dog and its handler) causing the two phenomena (high performances during training and validation). Characteristics could include a genetic factor enabling the dog to easily detect SARS-CoV-2 [41] or a good human-dog relationship within the dyad [42]. This hypothesis is supported by the heterogeneous performances during training and validation despite identical working hours for all dogs and by the good performances of the two “green” dogs (Ace and Bolt) during training (89% and 94%, respectively) and validation (86% and 100%). If a true causal association is causing the significant correlation, then increasing the dog’s performance during training is likely to increase the performance during validation. If increasing the number of training days increases training performances, then one reason for the heterogenous overall sensitivities in our study might be that the dogs with the lowest sensitivities required more than two weeks training. Our data did not enable the statistical association between performances during training and validation to be estimated after taking potential confounders into account.

The two week training period was much shorter than reported in some detection dog studies, but was in accordance with two previous studies on COVID-19 detection dogs [2, 3]. The training (Table 3) and validation (Table 6) performances support the hypothesis that two weeks of intensive training seems sufficient for most explosives and cadaver detection dogs. Interestingly, one of the two “green” dogs reached an overall sensitivity of 100% and the other reached 86% suggesting that even dogs with no training in olfactory detection may be trained to detect COVID-19 positive individuals. Further studies identifying factors associated with elevated training and validation performances are necessary to maximize training efficacy.

The main limitation of our study was the absence of data on dogs marking COVID-19 negative samples. This prevented us from calculating lower and higher bounds of estimated sensitivities, specificities, and positive and negative predictive values, as recommended for detection dogs studies.[43] Therefore, we cannot rule out that the dogs had a high sensitivity with poor specificity. Although low specificity values are possible, it is not expected based on the training performances. Since around 70% of the samples used in the training line-ups (step 4) were COVID-19 negative, it would not be possible to obtain a low specificity if the sensitivity and global performance during training were high. For example, for the five dogs reaching an overall sensitivity of 100% (Allo, Bolt, Cody, Nox and Ramos) during validation, the lowest performance in step 4 of training was 86% (Nox). If Nox’s sensitivity had been 100% in training, Nox’s specificity would have to be 79% to reach a global performance of 86%. Further studies collecting data for COVID-19 positive and negative samples are necessary to estimate sensitivities and specificities, and to calculate positive and negative predictive values.

Since the beginning of the COVID-19 pandemic, RT-PCR has been the reference testing method, and is considered reliable for SARS-CoV-2. However, Axell-House et al. recently evaluated the diagnostic accuracy of the COVID-19 test in a scoping review [44]. They found substantial heterogeneity among available studies in terms of test types, reference standards, metrics, and details of study design and methodology, and concluded that “while more than 200 SARS-CoV-2 molecular diagnostic tests have received FDA [Emergency Use Authorizations], we have described […] that the performance of few of these tests has been assessed appropriately. […] The accuracy of these tests should be interpreted with caution”. In this context, the sensitivities of the 21 study dogs may have been underestimated or overestimated since their calculation was based on the fact that the RT-PCR is a “gold standard” with a positive predictive value of 100% (i.e. the 151 COVID-19 positive patients were all truly infected by SARS-CoV-2) [45].

Twelve dogs always had 5-cone line-ups. For the remaining dogs, a maximum of 10% of line-ups had seven cones (Table 6). Cone number had little effect on overall sensitivities. Looking just at the 5-cone line-ups containing four or five samples, the overall sensitivities were lower for five dogs, remained the same for 5 other dogs, and was increased for the 11 remaining dogs (data not shown). Line-ups with few samples had the benefit of being easier to “match” since matching line-ups with one positive and one negative sample was easier than one positive and three or four negative samples.

In detection dog studies using line-ups, to be confident that dogs detect the studied disease itself and not a characteristic specific to the disease (a confounding factor), disease and disease-free samples in the line-up must be comparable, except for the disease status [34, 39]. As with observational studies (such as case-control studies), matching disease and disease-free samples in a line-up for more than one or two potential confounding factors would have been complicated. This is because the sweat sample had to be sniffed within days after sampling making it impossible to wait for an appropriate matched sample. Furthermore, the daily number of SARS-CoV-2 infected patients presenting in one hospital who undergo a RT-PCR test was not high enough to find matching COVID-19 negative patients from the same hospital. The hospital was the only characteristic which could have been matched for COVID-19 negative and positive samples but was not possible in the present study for organisational reasons. Apart from this characteristic, matching potential confounding factors in detection dogs studies working with line-ups should be performed a posteriori, when performing statistical analyses.

“Matched” sensitivities were calculated after the overall sensitivity for each dog. Theoretically, if a characteristic was a confounding factor causing high overall sensitivities, matching for this characteristic would have systematically decreased sensitivity in all 21 dogs. This decreased sensitivity was only observed for eight of the 21 dogs when matching for anosmia and asthma (Table 7). Most of the time, sensitivities increased after matching. Hypertension, dry cough, and sore throat were not considered potential confounding factors. However, sensitivity decreased for eight dogs when matching for hypertension and sore throat and for seven dogs when matching for dry cough (data not shown). Overall, these results do not prove that the overall sensitivities were due to a specific characteristic, and not SARS-CoV-2.

Some characteristics, other than the ones collected, could possibly cause a residual confounding bias such as medication use. However, matching for health conditions was likely to remove any confounding bias, at least for medications related to this condition, but it cannot be ruled out that other medications could be a confounder. Future studies should collect data regarding medication use for matching.

Our study has some strengths in accordance with previous recommendations for detection dog studies [33–35]. These include different samples being used in the training and validation sessions, randomised sample position, a large number of study dogs (of the 54 detection dog studies reviewed by Johnen et al. in 2017, only three involved more than 21 dogs[35]), only one sweat sample per individual and a large number of recruited individuals (n=261). Our study can be considered double-blind since the dog, its handler, and the data recorder were blinded to the sample locations. However, since the dedicated sample placement person remained in the room during the sessions, albeit with no visual contact with the dog or handler, the study cannot be considered fully blinded and therefore does not completely meet this recommended criterion. Despite the lack of visual contact, some unintentional influence on sample choice cannot be totally ruled out.

## CONCLUSION

A total of 1786 trials were performed during the training period, and 1368 during the validation period (using 151 positive and 110 negative samples). Calculated sensitivities range from 71% to 100%, but only three of the 21 dogs were under 80%, allowing the U.A.E. authorities to deploy the dogs in three international airports.

These results show that major conditions such as diabetes do not interfere with olfactory detection of COVID-19 in sweat, but confirmation is required with more specific studies. Future studies will focus on comparative field-test results including the impact of the main COVID-19 comorbidities and other respiratory tract infections and comparison with the different rapid COVID-19 tests in use.

## Notes

### Competing Interest Statement

The authors have declared no competing interest.

